# Causal modulation of right fronto-parietal phase synchrony with Transcranial Magnetic Stimulation during a conscious visual detection task

**DOI:** 10.1101/859330

**Authors:** Chloé Stengel, Marine Vernet, Julià L. Amengual, Antoni Valero-Cabré

## Abstract

Correlational evidence in non-human primates has reported evidence of increased frontoparietal high-beta band (22-30 Hz) synchrony during the endogenous allocation of visuospatial attention. But may the engagement of inter-regional synchrony at this specific frequency band provide the causal mechanism by which top-down processes are engaged and they facilitate visual perception in humans? Here we further analyzed electroencephalographic (EEG) signals from a group of healthy human participants who performed a conscious visual detection task, under the influence of brief *rhythmic* (30 Hz) or *random* bursts of Transcranial Magnetic Stimulation (TMS), with an identical number of pulses and duration, delivered to the right Frontal Eye Field (FEF) prior to the onset of a lateralized near-threshold target. We report increases of inter-regional synchronization in the high-beta band (25-35 Hz) between the electrode closest to the stimulated region (right FEF) and parietal leads, and increases of local inter-trial coherence in the same frequency band over parietal contacts, both driven by *rhythmic* but not *random* TMS patterns. Importantly, such increases were accompained by increases of visual sensitivity for left visual targets (contralateral to the stimulation) in the *rhythmic* but not the *random* TMS condition at the group level. These outcomes suggest that human high-beta synchrony between right frontal and parietal regions can be modulated non-invasively and that high-beta oscillatory activity across the right dorsal fronto-parietal network may contribute to the facilitation of conscious visual detection. Our study also supports future applications of non-invasive brain stimulation technologies for the manipulation of inter-regional synchrony, which could be administered to improve visual behaviors in healthy humans or neurological patients.

## Introduction

High-level cognitive functions, such as spatial attention orienting or access to perceptual consciousness, cannot solely rely on the activity of single cortical regions but require the integration of processes occurring in widely distributed cortical nodes organized in complex networks (Buzsáki & Draguhn, 2004; Varela et al., 2001). Accordingly, understanding how distant regions communicate as part of a single distributed network during the performance of a cognitive task has become a crucial mission for the systems neuroscience field.

Early theories of inter-regional communication in the human brain focused on the study of anatomical white matter connectivity, considered to be the backbone of long-distance communication (Laughlin & Sejnowski, 2003; Mesulam, 1990). However, neuronal activity and patterns of functional connectivity show episodic dynamic fluctuations operating in the order of milliseconds (Bressler & Tognoli, 2006; Britz et al., 2010). Consequently, these cannot be conveyed solely by the architecture of inter-areal structural pathways (a.k.a., the connectome), which is lacking the flexibility for a dynamic and selective communication between subsets of brain systems organized as nodes within highly interconnected networks (Fries, 2005).

For the last two decades, new mechanistic models have proposed that local and interregional communication between neuronal populations is subtended by the synchronicity of their oscillatory activity (Engel et al., 2001; Fries, 2005, 2009; Fries et al., 2001; Varela et al., 2001). Such models have claimed that when two natural cortical oscillators synchronize in frequency and/or in phase, the spikes generated by a first group of neurons will reach well-synchronized neurons within a target population at their peak of excitability, ensuring a higher gain in information transfer and, consequently, more efficient communication. This so called model of *communication-through-coherence* (Fries, 2005; Fries et al., 2001) has been hypothesized as particularly important in mediating top-down modulation (e.g. by attentional or perceptual processes) of inputs signals entering primary sensory areas (Engel et al., 2001).

Experimental data in support of long-distance synchronization during visual perception and the orientating of attention have been collected both in animal models (Buschman & Miller, 2007; Gregoriou et al., 2009; Saalmann et al., 2007) and humans (Gross et al., 2004; Hipp et al., 2011; Rodriguez et al., 1999). The former evidence suggests that in such processes, fronto-parietal regions may synchronize at a beta or gamma frequency bands (ranging from 15 to 60 Hz) during episodes of attentional orienting or perception. However, these reports also associated synchronization with specific behaviors solely on the basis of their co-occurrence in time, and have proven unable to distinguish causal contributions of oscillatory activity from potential epiphenomena.

The use of Transcranial Magnetic Stimulation (TMS), a non-invasive technology to stimulate circumscribed cortical regions, in combination with Electroencephalography (EEG), offers a unique approach to probe the causal implication of oscillations and synchronization patterns between cortical locations in behavioral effects likely encoded through such mechanisms. Indeed, TMS has demonstrated the ability to non-invasively manipulate attentional and perceptual behaviors (see examples in Chanes et al., 2012, 2013; Klimesch et al., 2003; Romei et al., 2010; Sauseng et al., 2009) as well as local and network activity correlates (Valero-Cabré et al., 2007, 2005, see Valero-Cabré et al., 2017 and Polanía et al., 2018 for recent reviews) likey by inducing, interfering or modulating ongoing neural signals from circumscribed cortical sites.

More recently, it has also been shown that the delivery of brief bursts of TMS pulses (usually 4-5 pulses) regularly spaced in time (a pattern called *rhythmic* TMS), structuring a regular alpha rhythm around ~10 Hz (i.e ~1 TMS pulse every 100 ms), progressively phase-locked natural alpha oscillators over the posterior parietal cortex in human brains not enganged in any specific task (Thut et al., 2011). Ensuing studies have reported evidence supporting the ability of rhythmic TMS, applied in a wide range of frequency bands, to modulate performance in different cognitive processes (Hanslmayr et al., 2014; Jaegle & Ro, 2014; Romei et al., 2011, 2010). Finally, a recently published report by our group, based on the same dataset we here aimed to further analyze, provded evidence that 30 Hz rhythmic TMS delivered over the right Frontal Eye Field (FEF), a key region of dorsal attention orienting networks, was able to locally entrain high beta oscillations in the frontal region below the stimulation coil. Most important, it suggested that local entrainment could be causally linked to improvements of conscious visual detection for left-lateralized near-threshold targets (Vernet, Stengel et al., 2019).

Through the above mentioned studies, rhythmic TMS has built a solid credibility as the only tool able to causally explore the oscillatory basis underlying the top-down modulation of conscious visual perception in humans by identifying performance shifts tied to the entrainment of local rhythmic activity at specific frequency bands and cortical sites. However, the role of the interregional synchrony between a stimulated region (in the current case the right FEF) and other areas of the attentional orienting networks and the ability of this biomarker to be involved in the modulation of conscious visual perception during the delivery of rhythmic TMS pulses remains unexplored.

To this end, given indirect evidence showing that TMS-evoked oscillations can spread through connections to distant regions (Rosanova et al., 2009), we further analyzed the data of an existing TMS-EEG dataset (Vernet, Stengel et al., 2019), aiming to extend prior results supporting a perceptual modulatory role of local episodic entrainment of high-beta activity in the right FEF. We hypothesized that brief entrainment of local 30 Hz oscillations by rhythmic TMS patterns on the right FEF would induce transient inter-regional phase synchronization at a high beta frequency, likely within a dorsal fronto-parietal system linking this frontal site with posterior parietal areas (Capotosto et al. 2009; Quentin et al. 2014, 2015). Such hypothesis would be substantiated in EEG recordings with higher 30 Hz phase-locking values between the frontal EEG contact closest to TMS stimulation site and parietal contacts for *rhythmic* compared to *random* TMS patterns. We also hypothesized that, through increased fronto-parietal phase synchronization, rhythmic stimulation over the right FEF would entrain high-beta oscillations in ipsilateral posterior parietal cortical sites. Substantiating these effects, we predicted increases of local high beta power and inter trial phase coherence (ITC) over right parietal electrodes during *rhythmic* compared to *random* TMS patterns.

Crucially, our study was based on a carefully designed control condition (a *random* TMS pattern) delivering the same number of TMS pulses, thus, the same amount of stimulation as a *rhythmic* 30Hz pattern of interest, but without a regular frequency-specific spacing of TMS pulses. As shown in prior publications, this strategy enabled us to isolate the effect of the rhythmicity of the stimulation on inter-regional synchronization as well as the entrainment of local oscillations (Chanes et al., 2013; Quentin et al., 2015; Vernet, Stengel et al., 2019).

## Material and Methods

### Participants

We performed further analyses on an existing TMS-EEG dataset used in a recent publication to demonstrate local high-beta entrainment during stimulation of the right FEF and its association with improvements of conscious visual performance (see Vernet, Stengel et al., 2019 for details). A group of right-handed 14 healthy participants (9 women) aged between 20 and 34 years old (24 ± 4) took part in the original experiment. Participants had normal or corrected-to-normal vision. They all took part voluntarily after having signed a consent form and were naïve as to the purpose of the experiment. All the experimental procedures were performed according to the Declaration of Helsinki. A research protocol including all the interventions of this study was sponsored by the INSERM and approved by an Institutional Review Board known in France as *Comité de Protection de Personnes* (CPP Ile-de-France IV).

### Conscious visual detection paradigm

An in-house MATLAB (Mathworks, version R2012b) script using the Psychtoolbox extensions (Brainard, 1997) was used to control the presentation of visual stimuli synchronized with the delivery of the TMS pulses. During the task, participants were seated with their heads resting on a chin-rest set so that their eye’s canthi remained 57 cm away from the center of the screen.

Each trial started with a gray resting screen lasting for 2.5 secs, followed by a fixation screen that displayed a central fixation cross (size 0.5×0.5°) and a right and left rectangular placeholders (6.0×5.5°) drawn 8.5° away from the center (Fig. 1A). These placeholders indicated the potential right or left lateralized locations of the target during the trial. The duration of the fixation screen was jittered between 1.0 and 1.5 secs to avoid predictability with regards to upcoming events. A brief-lasting (66 ms) size increase (0.7 × 0.7°) of the central fixation cross alerted participants of the presentation of an upcoming target. After an inter-stimulus interval of 233 ms, in 80% of the trials a target appeared in the middle of the left or the right placeholder with equal probability. The remaining 20% of the trials were ‘catch’ trials in which no target was shown in any of the placeholders. The target consisted of a low-contrast Gabor stimulus (0.5°/cycle sinusoidal spatial frequency, 0.6° exponential standard deviation) with vertical lines, appearing for 33 ms. Stimulus contrast was individually adjusted for each participant during a calibration block carried out prior to the onset of the experimental session. At all times, contrast level was never set below 0.005 Michelson contrast units. Similar tasks had been employed in prior publications in our research group (see Chanes et al., 2013, 2015; Quentin et al., 2015; Vernet, Stengel et al., 2019).

**Figure 1.**
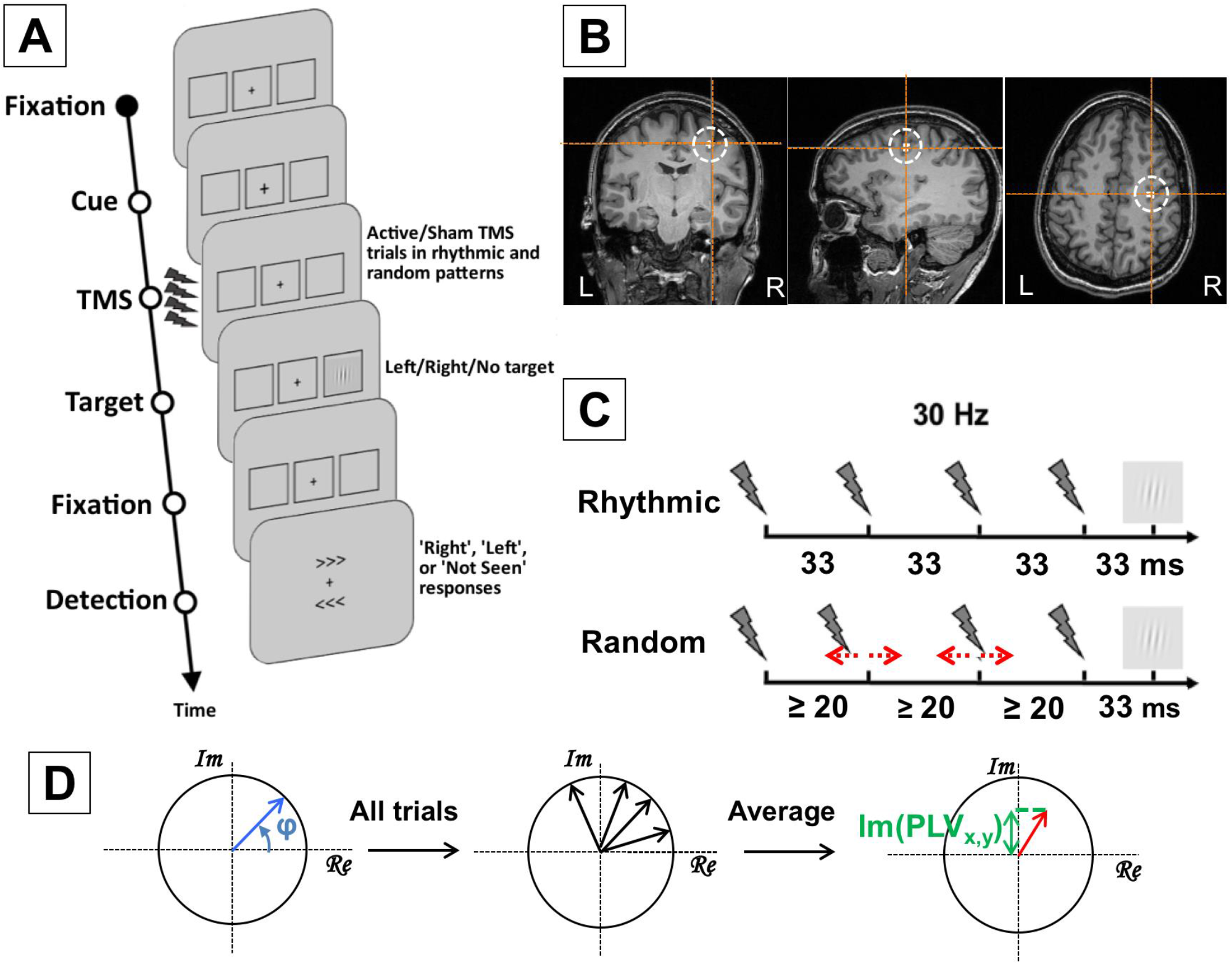
Experimental design, stimulation patterns, targeted cortical regions and Synchrony measures. **(A)** Visual detection task performed by participants. After a period of fixation, a central cross became slightly larger to alert participants of an upcoming event; then active and sham *rhythmic* or *random* TMS patterns were delivered to the right FEF region prior to the presentation of a target that could appear at the center of a right or left placeholder for a brief period of time. Participants were requested to indicate whether they did perceive a target or not and, if they saw it, where it appeared (right/left). Notice that in 20% of the trials, no target was presented in any of the placeholders. **(B)** Coronal, axial and sagittal MRI sections from the frameless stereotaxic neuronavigation system showing the localization of the targeted cortical right FEF (Talairach coordinates x=31, y=-2, z=47) in a T1-3D MRI of a representative participant. **(C)** Schematic representation of the TMS patterns employed in active and sham 30 Hz *rhythmic* stimulation to entrain oscillatory activity at the input frequency in the right FEF, and the *random* stimulation used as a control to isolate the effect of stimulation frequency. **(D)** Schematic representation of the calculation of the phase locking value. The phase of signals between each two sets of electrodes was extracted from the Fourier spectra. For all trials, complex vectors were reconstructed with a phase (φ) equal to the phase difference between two signals. These complex vectors were averaged over trials. TMS volume conduction was ruled out by calculating the imaginary phase locking value, which consisted in the projection on the imaginary axis of the complex vector averaged over all trials.

Participants were asked to perform a detection task, in which they had to report if they consciously saw the target and, if ‘yes’, to report on which side it appeared (‘right’ or ‘left’). The response window consisted in two arrow-like signs (“>>>“ and “<<<“) presented simultaneously below and above the fixation cross signaling the location of the right and left rectangular placeholders. Participants were requested to indicate which arrow pointed to the location of the placeholder where they had seen the target. The location of each arrow was randomized across trials to prevent the preparation of a motor response prior to the appearance of the response window and to make sure that visual processing activities in the FEF, located close to the motor areas, were temporally segreated from motor decisions and finger responses. Participants provided a response using three keyboard keys: an upper key to indicate the upper arrow (corresponding to the ‘d’ letter key), a lower key to indicate the lower arrow (corresponding to the ‘c’ letter key) and the space bar to indicate that no target had been consciously perceived.

Participants performed 6 blocks: 1 calibration block, 1 training block and 4 experimental blocks (2 blocks for each TMS pattern: *random* or *rhythmic*, see details on TMS patterns below). The order of the experimental blocks was counterbalanced across participants. Each block was divided into sub-blocks of 20 trials. The length of the calibration and training blocks was variable, as the termination of these two blocks depended on individual performance. Experimental blocks consisted of 7 sub-blocks and lasted approximately 20 minutes each.

During the calibration block, target contrast was adjusted to reach a performance of 50% correct detections using a staircase procedure (Cornsweet, 1962). Initially, the Gabor contrast was set very high (Michelson contrast of 1). Then, on each trial, contrast increased or decreased according to the response of the participant. The initial step in contrast was equal to the initial contrast level (note that, regardless, contrast was always kept higher than 0.005 Michelson contrast). On each reversal, contrast steps were divided by two. When, in five consecutive trials, contrast varied by less than 0.01 Michelson contrast, we considered that the 50% detection threshold had been reached. The threshold was measured a second time using the same procedure. The two thresholds were then compared. If they differed by less than 0.01 Michelson contrast, the calibration block was terminated and the contrast used during the following blocks was the average between the two thresholds. If they varied by more than 0.01 Michelson contrast, the threshold was measured again. During the calibration block, only sham TMS patterns were delivered on participants’ right frontal cortex. At the end of each sub-block, participants were invited to take a short break.

Before starting the experimental blocks, participants underwent a training block during which trials with active TMS (see below for further detail on TMS procedure) were introduced. For all conditions (no target present, target on the right, target on the left) half the trials delivered active TMS wheread the other half delivered sham TMS patterns. The order of presentation of active and sham TMS was randomized for each sub-block of 20 trials. Participants’ performance during the training block was verified to ensure that it remained stable even with the intermixed active TMS trials. At the end of each sub-block in this training period, participants were alerted if their false alarm rate was higher than 50% and received feedback on their percentage of incorrectly reported target position and on their percentage of incorrect fixation. Between sub-blocks, the experimenter could also manually adjust the contrast. Once participants reached a stable performance, the experimenter decided to end the training blocks and start the experimental blocks. Experimental blocks were identical to training blocks (with the same feedback for the participants) except that target contrast was kept constant for all sub-blocks and participants were allowed to take a short break only every two sub-blocks.

### Recording of eye movements

During all blocks, the position of both eyes was monitored on each participant with a remote eye tracking system (Eyelink 1000, SR Research, at sampling rate of 1000 Hz). If the location of the participant’s eyes was recorded more than 2° away from the center of the fixation cross at any time between the appearance of the alerting cue and target offset, the trial was considered as non-fixated. The trial was re-randomized amongst the remaining trials in the sub-block and repeated. Non-correctly fixated trials were excluded from any data analysis.

### TMS procedure

TMS was triggered in synchronization with the presentation of the visual stimuli via a high temporal resolution multichannel synchronization device (Master 8, A.M.P.I.) connected to two biphasic repetitive stimulators (SuperRapid, Magstim) each attached to a standard 70 mm diameter figure-of-eight TMS coil. The coil in charge of delivering active TMS patterns was held tangentially to the skull above the location of the right FEF, with its handle oriented ~ parallel to the central sulcus at a ~45° angle in a rostral to caudal, lateral to medial direction. The other coil, which delivered sham TMS stimulation was placed close the stimulation site but positioned perpendicular to the skull directing the magnetic field away from the brain. The sham coil produced the same clicking noise characterizing the delivery of an active TMS pulse but did not project any active hence effective stimulation to the targeted right frontal cortex.

During the whole experiment, the position of the active TMS coil was tracked using a neuronavigation system (Brainsight, Rogue Research). A T1-weighted MRI scan (3T Siemens MPRAGE, flip angle=9, TR=2300 ms, TE=4.18 ms, slice thickness=1mm) was acquired for each participant and the right FEF was localized on each individual scan as a spherical region of interest of radius 0.5 cm centered on the Talairach coordinates x=31, y=-2, z=47 (Paus, 1996) (Fig 1B). Using this neuronavigation system the active TMS coil was kept within a ± 3mm radius from the center of the targeted site during the whole experimental session.

As done previously (Chanes et al., 2013; Quentin et al., 2015; Vernet, Stengel et al., 2019), the two types of TMS patterns employed in this experiment consisted in a burst of four TMS pulses, lasting 100 ms and ending 33 ms before the onset of the visual target. Two types of patterns were tested: a *rhythmic* pattern for which the pulses were delivered regularly at a frequency of 30 Hz (33 ms of inter-pulse interval within the burst) and a *random* pattern engineered not to deliver any specific or pure single frequency. In the *random* pattern, the time interval between the 1st and the 4th pulse within the burst were kept identical to those in the rhythmic pattern. However, the 2nd and 3rd pulses were randomly jittered below and above their exact onset timings in the *rhythmic* pattern (Fig. 1C). Some constraints applied to pulse timing randomization. First, since the time needed by the TMS capacitors to fully re-charge before delivering the next pulse was limited, two pulses could not be delivered less than 20 ms apart. Second, the onset time of the two middle pulses (the 2nd and the 3rd) had to be shifted at least 3 ms away from the timings of the same pulses in the *rhythmic* pattern, to ensure that *random* patterns would never deliver a perfectly regular 30 Hz frequency.

TMS stimulation intensity was set at a fixed level of 55% of the maximal simulator output for all participants. This level was slightly higher than the intensity proven efficient in prior studies by our team (Chanes et al., 2013, 2015; Quentin et al., 2014) to take into account the increased distance between the coil and the cortex due to the presence of the EEG electrodes and the cap. To allow across-study comparisons, at the end of the experiment, the individual resting motor threshold (RMT) in the right hemisphere was determined for each participant as the TMS intensity that yielded a motor activation of the *abductor pollicis brevis* muslce (thumb motion) in about 50% of the attempts (Rossini et al., 2015). On average, the RMT was at 72 ± 9% of maximum stimulator output. The fixed stimulation intensity (55% of maximal machine output) used for TMS corresponded to 78 ± 12% of our participant’s individual motor thresholds.

### EEG recordings

EEG signals were continuously recorded during all experimental blocks with a TMS-compatible system (BrainAmp DC and BrainVision Recording Software, BrainProducts GmbH). We recorded signals from 60 electrodes spread evenly across the scalp and positioned according to the international 10-20 system, plus a reference on the tip of the nose, a ground on the left ear lobe and 4 additional EOG electrodes positioned above and below the right eye and on each temple. Skin/electrode impedances were maintained below 5 kOhm. The signal was digitized at a sampling rate of 5000 Hz.

#### EEG epoching and artifact removal procedure

All EEG data analyses were performed with the FieldTrip toolbox (Oostenveld et al., 2010) running on MATLAB R2017b. The EEG and EOG data were epoched across a [-2, 2] seconds window centered on the onset of the target. Trials where the participant did not correctly fixate the central cross in the time interval between the alerting cue onset and the target offset were automatically excluded during the task performance by monitoring gaze position using an eye tracking system. Prior to any data analysis, all trials contaminated by blinks in the [-500 500] ms epoch around visual target onset were removed by visual inspection. After these procedures, an average of 126±13 trials remained for each experimental block.

To remove the artifact created by the discharge of a TMS pulse, data in a window of [-4, +12] ms centered on the delivery of a pulse was discarded. A second order Butterworth filter (1 to 50Hz), with forward-backwards filtering, was applied on the remaining data before the the gap of removed EEG data was filled with a piecewise cubic spline interpolation. To reduce the burden of computation time, data were down-sampled to 500 Hz before an Independent Component Analysis (ICA) was applied. To make sure that the ICA did not introduce any difference between conditions, trials for all 4 experimental blocks (whether they were active or sham TMS trials, and whatever the TMS pattern tested) were gathered together and the ICA was computed on this single dataset. This procedure enabled the removal of residual artifacts (including eye movements, electrode malfunctions, 50 Hz power line artifacts and TMS artifacts lasting longer than 12 ms). Components were identified as artifacts based on the guidelines of Rogasch et al. (2014). On average 9 ± 2 components were rejected. After this procedure, data was separated into 4 conditions: Active *rhythmic* TMS, Sham *rhythmic* TMS, Active *random* TMS and Sham *random* TMS.

#### TMS-EEG data analyses

Epoched EEG signals were transformed into the time-frequency domain using a 3-cycle Morlet wavelet transform on the time window [-500 +500] ms around target onset and for frequencies between 6 and 50 Hz. Three measures associatd to the notion of oscillation synchronization were calculated: power, inter-trial coherence (ITC) and imaginary phase-locking value (PLV). Power was calculated as the squared value of the modulus of the Morlet coefficients (per each time frame and frequency bin) relative to a baseline window [−500 −300] ms before target onset. This outcome measure shows, in decibels (dB) unit, the increase (positive value) or decrease (negative value) of power relative to this period.

Inter-trial coherence (ITC) estimates phase consistency across trials in a given location (an electrode or a group of electrodes). To measure synchronization between distant regions, we calculated the phase-locking value (PLV) (Lachaux et al., 1999). This measure reflects the stability of the phase difference between two signals in a specific frequency band. The phase difference between two signals is extracted at all points in time from their Fourier spectra and are then averaged across all trials. The PLV is defined as the module of the resulting averaged complex vector across trials (Fig. 1D). The following formula was used for its computation:

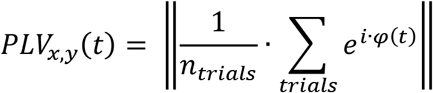

where *φ* (*t*) represents the phase difference between the signal recorded by the electrodes *x* and *y* at time *t* and *n_trials_* is the total number of trials in the condition.

The PLV, as the module of a unit vector, is always comprised between 0 (random phase difference between the two signals) and 1 (constant phase difference between two signals) (Guevara & Corsi-Cabrera, 1996). To avoid controversy on the fact that the PLV might be very sensible to volume conduction, which would bias this parameter to show higher values of phase synchrony over the whole scalp but particularly between neighboring electrodes (Vinck et al., 2011), we considered as a synchrony value the projection on the imaginary axis of the complex vector of the PLV, hence the so called *imaginary PLV* (Nolte et al., 2004; Vinck et al., 2011).

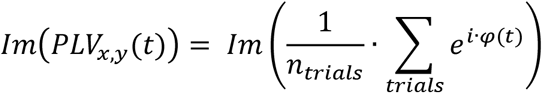

This value of synchrony effictively cancels the contribution to the PLV of signals with a null or close to null phase difference, which is characteristic of two highly correlated signals affected by volume conduction. The normalized value of the imaginary PLV is comprised between −1 (when the two signals compared have a constant phase difference of – π/2) and 1 (when the two signals compared have a constant phase difference of π/2). To reject any information about which signal in the pair was lagging in phase behind the other, we calculated the absolute value of the imaginary PLV (comprised between 0 and 1) and retained it to estimate synchronization. Such phase information is difficult to interpret due to the cyclicity of the phase and would only render our results more difficult to understand.

In our analysis, we computed the imaginary PLV between the electrode FC2 (the closest to the stimulation coil targeting the right FEF) and all other scalp EEG leads for the time window [−500, +500] ms centered around visual target onset and for frequencies between 6 and 50 Hz.

#### Behavioral Data Analysis

Performance in the detection task was assessed through the perceptual sensitivity index (*d’*), a measure from Signal Detection Theory that quantifies objective perception of stimuli presented around the threshold of perception (Stanislaw & Todorov, 1999). Trials were separated in ‘hits’ (when the target was correctly detected), ‘misses’ (when the target was not reported), ‘false alarms’ (when a target was reported for a catch trial, i.e. trials when no target was presented), ‘correct rejections’ (when no target was reported in a catch trial) and ‘errors’ (when a present target was reported on the wrong side of the screen). The perceptual sensitivity index was then calculated from the rate of ‘hits’ and ‘false alarms’. See Vernet, Stengel et al. (2019) for full details on such analyses.

#### Statistical analyses

We applied a 2×2 orthogonal design with TMS pattern (*rhythmic, random*) and TMS condition (*active*, *sham*). Therefore, values of imaginary PLV, power and ITC were compared in two different ways. First, for each TMS pattern (*rhythmic* and *random*) we contrasted the *active* TMS condition and the *sham* TMS condition. Second, for each stimulation condition (*active* and *sham*), we compared the *rhythmic* TMS and *random* TMS bursts. Each pair were compared with two-tailed paired Student’s *t*-test (*α* = 0.01). To correct for multiple comparisons in both topographical and time-frequency maps, we performed cluster-based permutation tests with Montecarlo sampling. This method clustered together neighboring electrodes or time-frequency points that reached significance in the paired t-test, using a single *t*-value per cluster. A nonparametric permutation test was then applied on these clusters (10000 permutations, alpha = 0.05) to determine which ones survived the correction for multiple comparisons. Cluster-based permutations is a highly sensitive method for correcting for multiple comparisons in EEG data because it is adapted to data highly correlated in space and time (i.e. an effect on the EEG signal is likely to be spread over adjacent EEG leads and consecutive time points) (Maris & Oostenveld, 2007). However, currently no consensus exists on how cluster-based permutations should be applied in factorial designs to evaluate interaction effects between multiple factors (Edgington & Onghena, 2007; Suckling & Bullmore, 2004). For this reason, and guided by our *a piori* predictions of a contrast between rhythmic and random TMS patterns to isolate the impact of rhythmic structure of pulses within TMS patterns on oscillatory activity, we computed direct pairwise comparisons between conditions.

We initially focused our analysis on EEG signals recorded during stimulation (time window [-133 0] ms, 0 being the target onset) and in the high-beta frequency band of interest ([25 35] Hz) and computed the topographical maps for PLV between electrode FC2 (closest to the targeted right FEF) and all other electrodes on the scalp. However, the localization of significant effects revealed by permutation tests on the topography are not very precise. Indeed, the building of clusters of electrodes might blur the effect over larger regions. Additionally, it must not be forgotten that the null hypothesis tested in the permutation test in order to control for multiple comparisons extends to the whole array of electrodes and therefore does not permit to conclude on the localization of any of the observed effects (Maris & Oostenveld, 2007). In order to investigate in further detail the spatial localization of the synchronization induced by *rhythmic* or *random* TMS we defined two separate regions of interest, one in the left and one in right hemisphere, including parietal and parietooccipital electrodes locations, and we compared time-frequency maps of PLV, Power and ITC for these two locations.

To analyze the participants’ performance, a 2×2×2 repeated measures analysis of variance (ANOVA) was applied to values of perceptual sensitivity index (d’) with *stimulation pattern* (rhythmic, random), *stimulation condition* (active, sham) and *target location* (left, right) as within-participant factors. Planned post-hoc t-student tests were also used for pairwise comparisons. See Vernet, Stengel et al. (2019) for more detail on this analysis.

## Results

### High-beta band right fronto-parietal synchronization

We illustrate (Fig. 2) the topographic distribution of imaginary PLV within the high-beta band (25-35 Hz) between contact FC2 (in the scalp EEG array, the closest to the targeted right FEF) and the remaining 59 scalp electrodes, during the TMS delivery period ([-133 0] ms). Statistical analyses revealed for such frequency band (around the stimulation frequency of rhythmic TMS patterns) and time window (TMS burst duration) statistically significant differences between the two types of active tested TMS patterns. More specifically, *rhythmic* active, compared to *random* active TMS patterns, increased synchronization between the frontal electrode FC2 and a group of leads overlying right parieto-occipital regions ipsilateral to the delivered stimulation. The same pattern of fronto-parietal synchronization was also observed when contrasting active *rhythmic* stimulation to sham TMS bursts.

**Figure 2.**
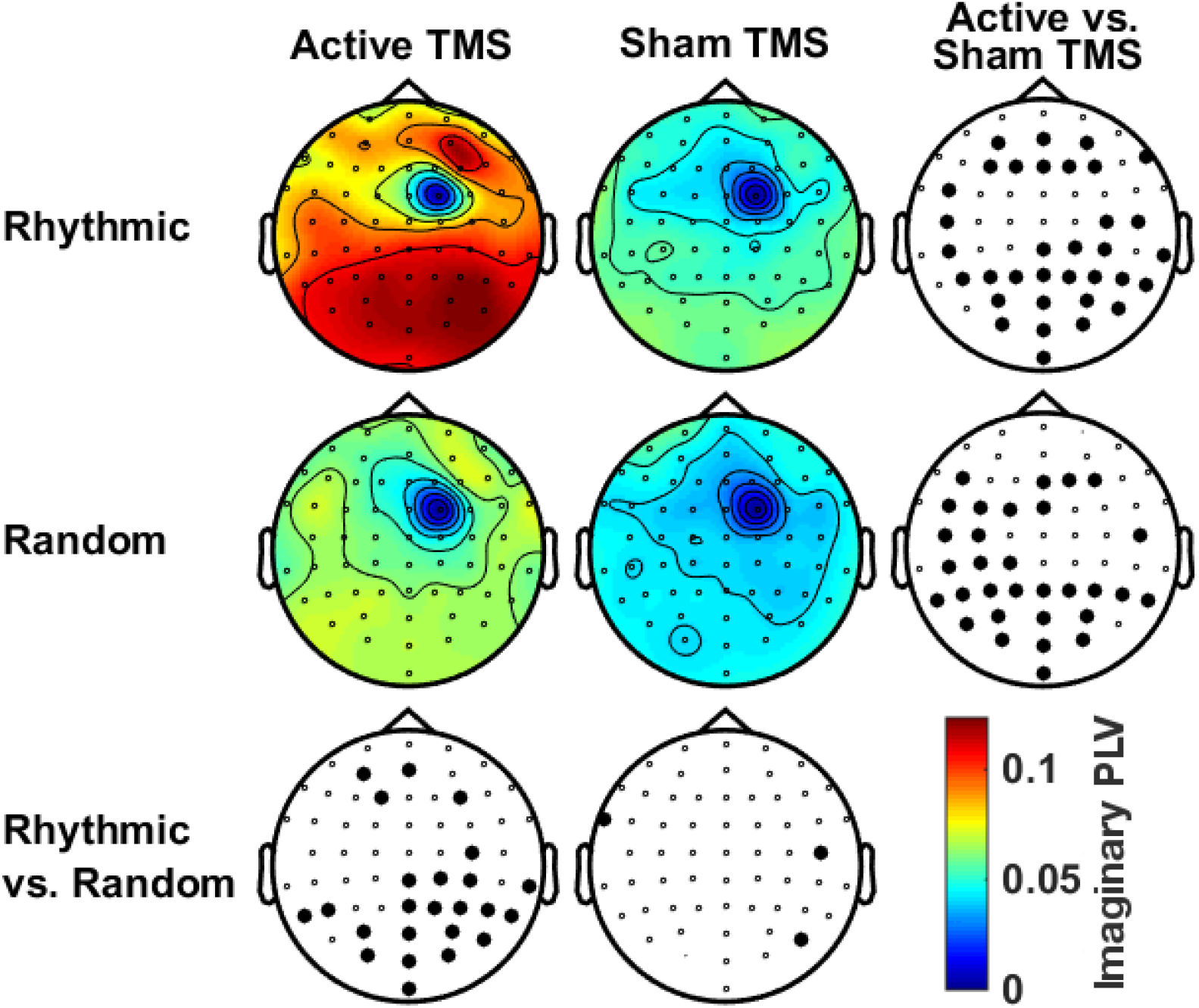
Topographical maps of high-beta imaginary PLV distribution conditions across stimulation patterns delivered to the right FEF. Topographic maps represent imaginary synchrony values (Imaginary PLV) for each scalp electrode compared to the signal recroded by the electrode closest to the targeted right FEF region (10-20 EEG system lead FC2) and the TMS coil center, for a time window preceding visual target onset [-0.133 0] ms and high beta frequency band [25 35] Hz. Maps are arranged following our 2×2 cross-design, comparing sham vs. active TMS condition (columns) and random vs. rhythmic TMS patterns (rows). Bottom and right topographies present the outcomes of the statistical permutation tests, with large black dots indicating EEG electrodes for which imaginary PLV differences reached statistical significance between TMS conditions (p<0.05), either active/sham rhythmic TMS vs. active/sham random TMS (bottom row) or active rhythmic/random TMS vs. sham rhythmic/random TMS (right column). Note that imaginary synchrony increased significantly in a group of right parietal electrodes for the active rhythmic TMS condition compared to both the sham rhythmic TMS and active random TMS control patterns.

Statistical analyses on topographic maps also suggest that *random* TMS patterns (comparison between *random* active and *random* sham conditions) increased synchronization around 30 Hz between the right FEF and fronto-parietal regions in the left hemisphere (contralateral to the stimulation). Hence, in an attempt to refine the spatial distribution of such increased frontoparietal synchronization observed in response to *rhythmic* or *random* TMS, we defined two regions of interest comprised of left or right parietal and parieto-occipital scalp EEG contacts. Statistical analyses on time-frequency datasets (Fig 3) confirmed that, compared to *random* TMS bursts, *rhythmic* TMS patterns increased right fronto-parietal synchrony only during the delivery of active stimulation and that such effects operated within a frequency band restricted to high-beta (24-45 Hz) oscillations (Fig. 3A). A similar result emerged when comparing the *rhythmic* active and sham conditions. However, contrary to what was suggested by our first topographic analysis, no statistically significat effects were found for active *random* TMS patterns on left fronto-parietal synchrony (Fig. 3B comparison between *random* active and sham conditions).

**Figure 3.**
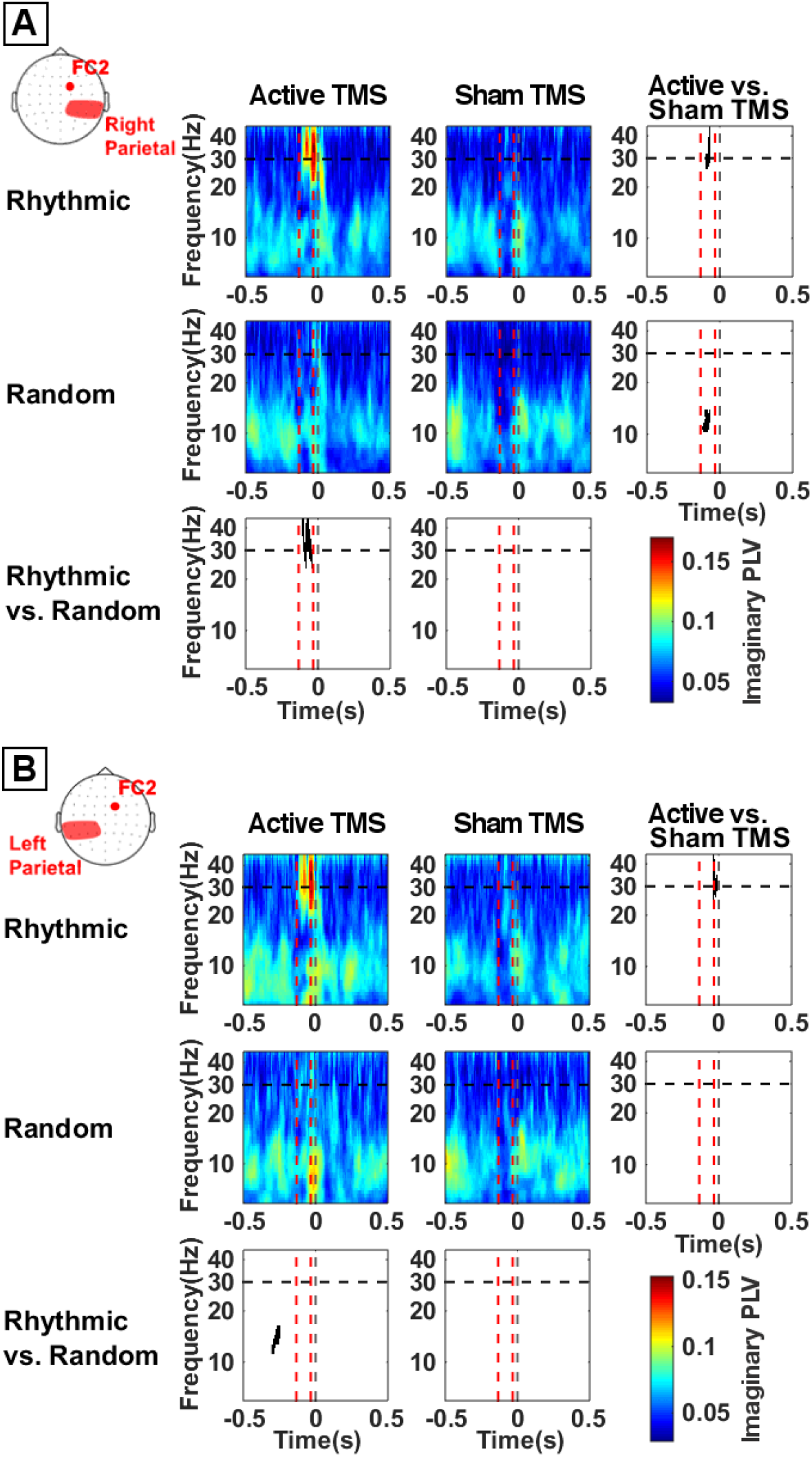
Temporal dynamics and frequency-specificity of fronto-parietal imaginary synchronization (PLV) across stimulation patterns delivered to the right FEF. Time frequency analyses of imaginary synchrony (Imaginary PLV) between the FC2 scalp EEG electrodes and right **(A)** or left **(B)** parietal scalp EEG leads. Time is centered (vertical dotted gray line, t=0) to the onset of the visual target. Panels are arranged following our 2×2 cross-design. We compared *sham* vs. *active* TMS condition (rows) and *random* vs. *rhythmic* TMS patterns (columns). Vertical dotted red lines signal the time window between the 1st (−133 ms) and the 4^th^ and last (−33 ms) TMS pulses of the delivered stimulation patterns. Horizontal dotted black line indicate the frequency of TMS rhythmic stimulation pattern (30 Hz). Bottom and right panels present the outcomes of the statistical permutation tests, with black pixels indicating clusters that reached statistical significance between TMS conditions (p<0.05) either active/sham rhythmic vs. active/sham random (bottom row) or active rhythmic/random vs. sham rhythmic/random (right column). Imaginary frontoparietal synchrony increased ipsilateraly (right hemisphere) during 30 Hz rhythmic stimulation compared to both random and sham TMS controls.

Topographic and time-frequency analyses on PLV between frontal and parietal scalp EEG contacts suggested that the delivery of *rhythmic* TMS patterns over the right FEF induced an increase of inter-regional synchronization in the high-beta band. Such increases proved short-lasting (i.e., not exceeding the period of the 4 pulse TMS burst) and likely entrained across a frontoparietal network of cortical sites within the targeted right cerebral hemisphere.

### Long-distance oscillatory parietal entrainment of high-beta band activty

Oscillation power within the 25-35 Hz high-beta band in both left and right regions of interest (i.e., either in parietal or in parieto-occipital EEG contacts) increased significantly in response to active *rhythmic* TMS patterns delivered frontally (compared to sham *rhythmic* TMS burst), but not during active *random* frontal stimulation (compared to its equivalent sham TMS bursts) (Fig. 4).

**Figure 4.**
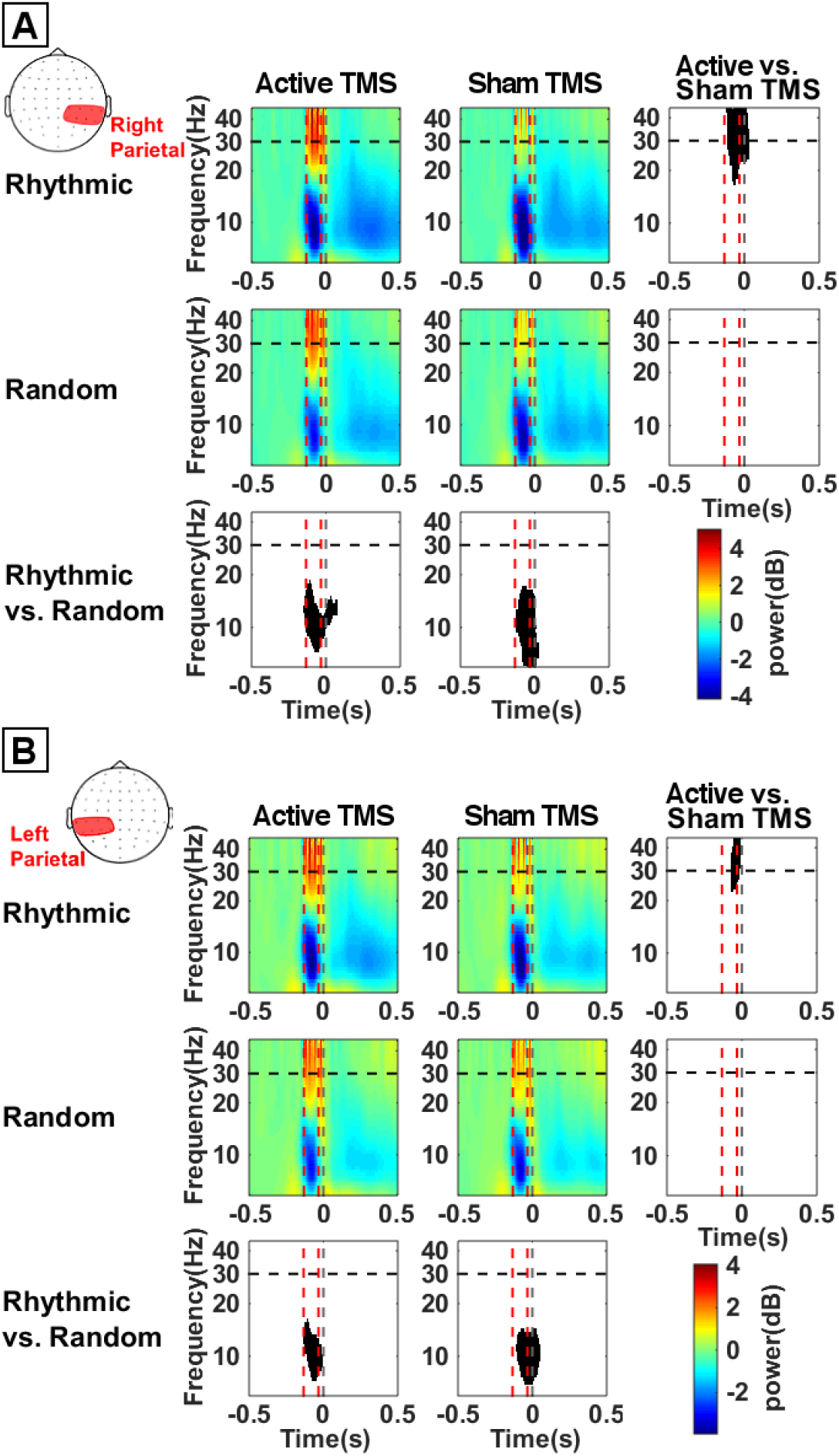
Temporal dynamics and frequency-specificity of high beta local entrainment (power) on parietal regions across stimulation patterns delivered to the right FEF. Time frequency plots representing oscillation power over right parietal **(A)** or left parietal **(B)** electrodes. Time frequency panels are arranged following a 2×2 cross-design, in which we compare *sham* vs. *active* TMS condition (rows) and *random* vs. *rhythmic* TMS patterns (columns). Time is centered on the onset of the visual target (vertical dotted gray line, t=0). Vertical dotted red lines indicate the time window between the 1st (−133 ms) and the 4^th^ and last (−33 ms) TMS pulses of the delivered stimulation pattern. Horizontal dotted black line indicates the delivered frequency of stimulation (30 Hz) for rhythmic patterns. Bottom and right panels present the outcomes of the statistical permutation tests, with black pixels indicating clusters that reached statistical significance between TMS conditions (p<0.05), either active/sham rhythmic vs active/sham random (bottom row) or active rhythmic/random vs. sham rhythmic/random (right column). Notice that right frontal rhythmic stimulation, compared to sham stmulation, transiently increased high-beta oscillation power over the left and right parietal cortices, hence at distance from the stimulation site. However, no differences were found for the time course of 30 Hz power comparing active rhythmic vs. active random TMS patterns. Note also that the sound of the TMS coil delivering rhythmic stimulation might be responsible for the decerase (desynchronizaton) of alpha oscillations in bilateral parietal regions.

Inter-trial coherence (ITC), a measure of phase-locking of local oscillations, was also found to be significantly increased in left and right parietal regions of interest following active rhythmic TMS to the right FEF (Fig. 5). Indeed, the comparison between active and sham TMS conditions showed higher ITC levels during active stimulation for both, *rhythmic* and *random* TMS patterns in bilateral parietal sites. However, a direct comparison between the two active conditions (*rhythmic* vs. *random* TMS patterns) revealed higher levels of a trial-by-trial phase-locking for *rhythmic* TMS patterns compared to *random* TMS patterns.

**Figure 5.**
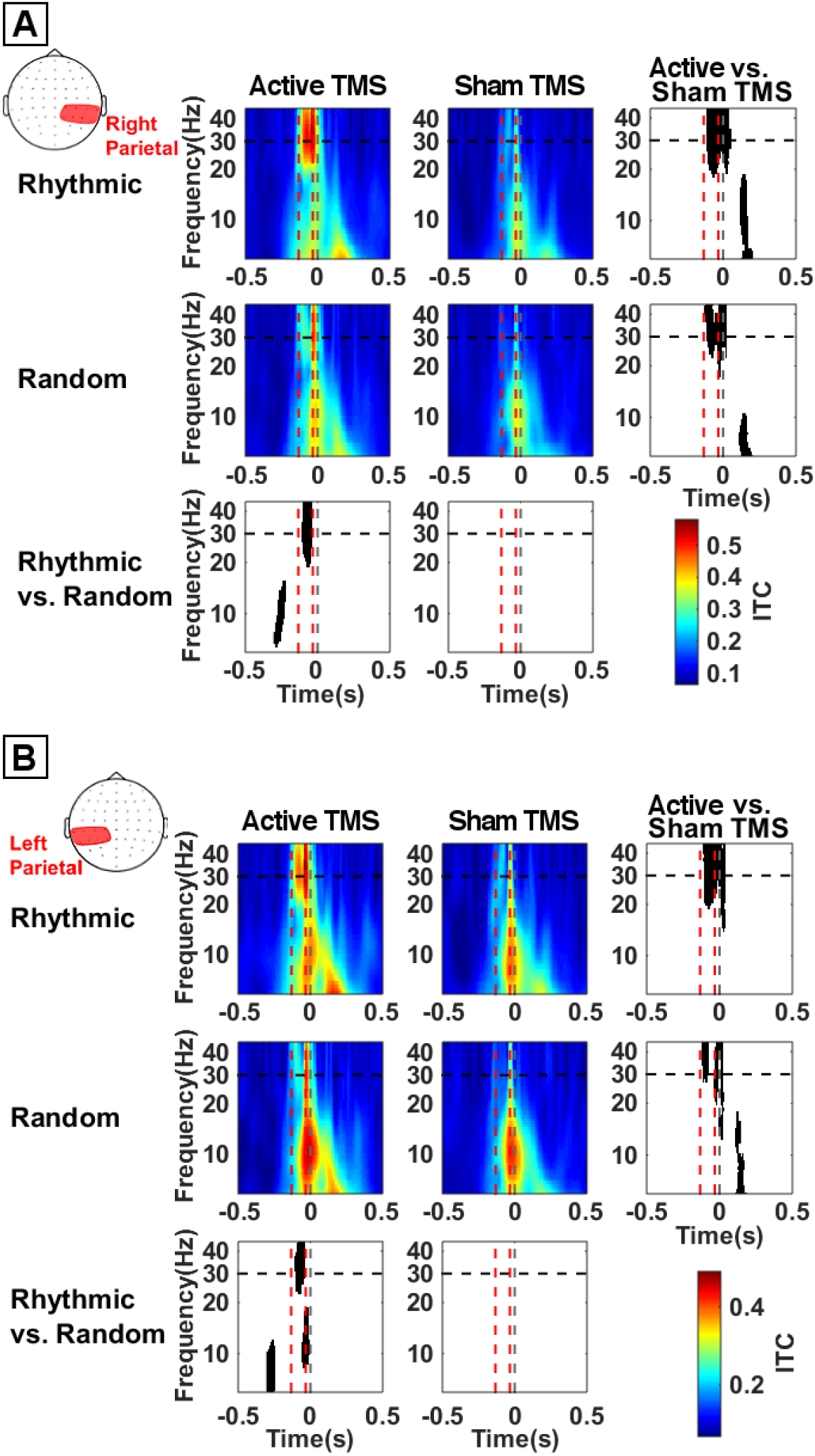
Temporal dynamics and frequency-specificity of distant phase-locking of high beta oscillations on parietal regions across stimulation patterns delivered to the right FEF. Time frequency plot of inter-trial coherence (ITC) over right parietal **(A)** and left **(B)** parietal scalp EEG electrodes. Time frequency panels are arranged following a 2×2 cross-design, in which we compared *sham* vs. *active* TMS condition (rows) and *random* vs. *rhythmic* TMS patterns (columns). Time is centered on the onset of the target (vertical dotted gray line, t=0). Vertical dotted red lines indicate the time window between the 1st (−133 ms) and the 4^th^ and last (−33 ms) TMS pulses of the delivered stimulation pattern. The horizontal dotted black line indicates the frequency of rhythmic TMS stimulation (30 Hz). Bottom and right panels present the outcomes of the statistical permutation tests, with black pixels indicating clusters that reached statistical significance between TMS conditions (p<0.05), either active/sham rhythmic vs active/sham random (bottom row) or active rhythmic/random vs. sham rhythmic/random (right column). Note that rhythmic stimulation of the right FEF increased inter-trial coherence in the high-beta range distantly over right and left parietal EEG contacts, whereas random stimulation patterns increased phase-locking transiently in the alpha frequencies in parietal EEG leads rather contralateral to right FEF stimulation.

### Ryhthmic and random TMS effects in alpha band synchronization

Aside from the modulation of oscillatory activity in the high-beta band, our data also unexpectedly revealed alpha desynchronization over parietal areas for all 4 TMS conditions (active or sham, rhythmc and random stimulation) during burst delivery (Fig. 4). This outcome might not be likely associated to TMS active fields, as the delivered patterns failed to show any significant difference when comparing active vs sham TMS patterns. However, both statistical *rhythmic*/*random* comparisons (active *rhythmic* vs. active *random* and also sham *rhythmic* vs. sham *random*) showed that *rhythmic* patterns induced stronger alpha desynchronization than *random* patterns.

By the end of the TMS burst, we observed alpha band phase-locking increases over the same parietal contacs (Fig. 5). Although visible in all 4 conditions (note that, again, this phaselocking was not significantly different between active and sham TMS patterns), phase-locking was this time stronger for active *random* stimulation compared to active *rhythmic* stimulation) in left parietal contacts (hence located on the hemisphere contralateral to the stimulation) (Fig. 5B). Further investigations with a more adapted behavioral paradign will be necessary to better pinpoint the origin and potential contribution of such alpha band activity.

### Effect of high-beta synchronization on conscious visual detection

As already reported (Vernet, Stengel et al., 2019), *rhythmic* stimulation at 30 Hz delivered on the right FEF modulated the outcomes of the detection task performed by particpants. A 2×2×2 ANOVA on perceptual sensitivity (d’) values revealed a main effect of *stimulation condition* (active or sham) F(1,13)=5.33; p < 0.05), with higher levels of visual sensitivity (d’), i.e. better detection performance, in trials in which the right FEF had been stimulated with active TMS, but indistinctively for rhythmic and random TMS patterns.

Statistical analyses also revealed a significant main effect of *visual field* (right, left) (F(1,13)=10.14; p < 0.01) with higher perceptual sensitivity (d’) for right targets compared to those presente in the left visual hemifield. No other significant effects were found, although a three-way interaction *visual field* x *stimulation pattern* x *stimulation condition* displayed a trend towards statistical significance (F(1,13)= 3.97; p < 0.068).

On the basis of a strong a priori hypothesis supporting different effects for *rhythmic* and *random* stimulation patterns on conscious visual perception (Chanes et al., 2013b, 2015; Quentin et al., 2015; and see Vernet, Stengel et al., 2019 for furter details on statistical analyses for visual outcomes), Student’s t-test were calculated to compare active vs. sham TMS patterns. These analyses revealed that *rhythmic* active TMS (compared to its equivalent sham TMS) increased perceptual sensitivity (d’) for targets displayed on the left visual field (p < 0.01) but not for those presented on the right (p>0.88) (Fig. 6). No significant differences between right and left targets were found between active *random* and sham *random* TMS patterns (both active vs. sham comparisons p>0.11).

**Figure 6.**
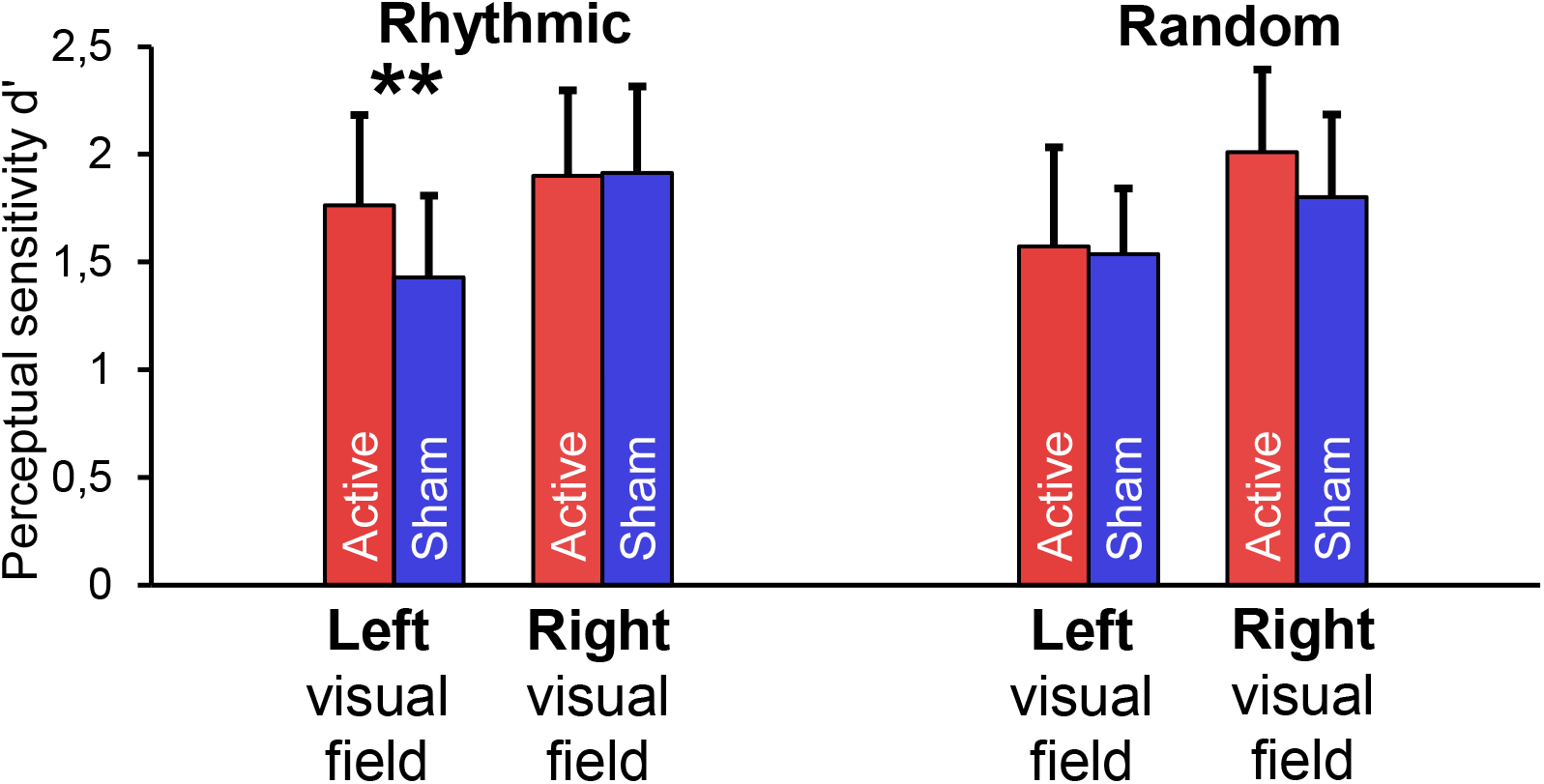
Modulation of perceptual sensitivity (d’) for left and right visual targets presented across stimulation patterns delivered to the right FEF. Data are presented for the rhythmic (Left) and random TMS stimulation conditions contrasting active (in red) vs sham (in blue) patterns for targets presented in the left (contralateral to right FEF stimulation) and the right (ipsilateral to right FEF stimulation) visual hemifield. For details on statistical approaches refer to the Results section. ** p < 0.01 in post-hoc t-tests between active vs sham conditions. Notice that perceptual sensitivity is increased only for targets contralateral to the stimulation site (left) in active compared to sham rhythmic TMS patterns. Decision criterion (not pictured) was not modulated by the stimulation. This behavioral outcomes on a conscous visual detection task are based in the same dataset previously presented (Vernet, Stengel et al., 2019).

## Discussion

Here we used rhythmic TMS coupled to EEG recordings during a conscious visual detection paradigm to explore in humans the contributions of 30 Hz inter-regional synchrony across right dorsal fronto-parietal systems and assess potential causal implications in the modulation of conscious visual perception. We demonstrated that high-beta rhythmic TMS patterns delivered over the right FEF induced synchronization in the high-beta band between this area and ipsilateral parietal regions. This increase in synchronization was transient and did not extend beyond the delivery of the last TMS pulse. Since synchronization increases were not observed when participants were stimulated with our control *random* TMS pattern with an equal number of pulses and total duration, we conclude that this effect is dependent on the specific spatio-temporal structure of our high-beta *rhythmic* pattern. We also posit that, very likely, such effects on frontoparietal synchrony are closely related to the local entrainment of a 30 Hz episodic rhythm induced by TMS in the right FEF, reported already in a previous analysis of this same dataset (Vernet, Stengel et al., 2019).

Given that focal stimulation has been shown to combine local and inter-regional effects (reviewed in Polanía et al., 2018; Valero-Cabré et al., 2017) we here investigated if a focal entrainment of a 30 Hz osicllation in frontal regions previously reported by our group (Vernet, Stengel et al., 2019) could be conveyed to distant interconnected cortical regions, not directly influenced by TMS patterns. To this end we focused on EEG leads overlying posterior parietal regions, notably those located in and around the intraparietal sulcus (IPS) which has been shown to interact with the right FEF as part of the dorsal attentional orienting network and shown an ability to modulate perceptual inputs (Capotosto et al. 2009; Quentin et al. 2014; Quentin et al. 2015).

As initially predicted, our analyses revaled that 30 Hz rhythmic TMS patterns delivered to the right FEF did not only entrain, as shown previously, right frontal high-beta activity (Vernet, Stengel et al., 2019) but increased high-beta power and phase-locked high-beta oscillations over parietal electrodes. This phenomenon can likely be explained by the entrainment of cortical oscillations via episodic TMS bursts delivered to a distant but richly interconnected area (see Thut et al., 2011). However, it is fair to notice that it could eventually be partially independent from direct intrahemispheric fronto-parietal synchronization mechanisms, as our data suggest that phase synchronization increased for only right EEG contacts but parietal entrainment was recorded for right and left parietal electrode contacts.

The electrophysiological effects induced by rhythmic TMS bursts delivered shortly before presentation of a low contrast target were accompanied by modulations of conscious visual performance, consisting in increases of perceptual sensitivity (d’) (Vernet, Stengel et al., 2019). Taken together, our results contribute evidence in favor of a causal implication of high-beta oscillatory activity, operating across fronto-parietal systems potentially involved in the allocation of spatial attention, in the modulation of conscious visual detection performance.

### Fronto-parietal brain rythms and the modulation of attention and perception

Our results are consistent with influential findings by Buschman and Miller (2007) in non-human primates, obtained by means of intracranial recordings and providing correlational evidence in favor of high-beta fronto-parietal synchrony during the allocation of endogenous attention in a top-down visual search task. Such work was replicated in humans by Phillips and Takeda (2009) employing scalp EEG. Our study used a simple conscious detection task which did not manipulate the allocation of attention by means of spatial or attentional cues. Nonetheless, this same modality of attention might have been engaged in our participants since, on each trial, they were cued with an central alerting central cue on the onset of an upcoming lateralized target with a given delay (~233 ms) for which only endogenous attention can uphold expectancy (Carrasco, 2011).

Our results build on the above-mentioned correlational results and, adding the value of causality, provide further proof in favor of a causal relationship between high-beta fronto-parietal synchrony and the modulation of visual perception in the human brain. Most importantly, they extend prior results derived from this same TMS-EEG dataset and suggest that short bursts of focal rhythmic TMS do not only show an ability to locally entrain frequency-specific rhythms within the targeted area dictated by the pace of stimulation (Vernet, Stengel et al., 2019) but can also be used to synchronize in a frequency-specific manner the stimulated target region (in our case the right FEF) with interconnected regions (such as posterior parietal sites) and entrain distantly rhythms at this same frequency.

Last but not least, our analyses also support the suitability of *rhythmic* TMS patterns (made as bursts of individually triggered pulses) delivered onto a specific cortical location and the comparison of their behavioral impact with those under the influence of equivalent control *random* patterns (of equal duration and number of pulses) allowing to isolate the causal contribution of oscilation frequency. Owing to the use TMS *random* patterns as a control condition, TMS-driven electrophysiological EEG differences between active *rhythmic* and *random* conditions here reported are unlikely to be artifacts caused by auditory (clicking) or tactile (scalp tapping) stimulation inherent to the delivery of TMS pulses. Neither could they be simply explained by the impact of a magnetic pulse on neuronal activity since potential artefactual effect of single pulses would be identically present in both active TMS conditions, *rhythmic* and *random*.

Pioneering research in this field has extensively employed rhythmic TMS (without coupled EEG recording) to investigate the causal role of local oscillatory activity on different cognitive processes subtending behavioral performance (Jaegle & Ro, 2014; Romei et al., 2010; Romei et al., 2011). Nonetheless, the current report is among the first to use coupled online TMS-EEG recordings and gather evidence supporting a causal link between a complex cognitive process such as the modulation of conscious perception subtended by long-range systems (such as dorsal attention orienting networks), specifically with modes of interregional synchronization between frontal and parietal locations.

### Causal role of inter-regional synchrony in visual perception improvements

The detailed mechanistic explanations for our electrophysiological effect and its associated behavioral findings remain open. Nonetheless in their original non-human primate study, Buschman and Miller (2007) hypothesized that synchronization of neuronal activity may increase the efficiency of inter-areal coordination and communication, enabling to process a single object and to suppress the processing of distractors. This hypothesis is consistent with the explanatory model developed in 2009 by Fries, in which inter-regional synchronization in the gamma-band was proposed to provide an exclusive and effective communication link between two areas selective for a visual target and invariant even in the presence of distractors (Fries, 2009).

Unfortunately, although our data show that entrained high-beta neural activity in the right FEF (modulated by focal rhythmic TMS bursts) increased synchronization between the stimulated site and electrodes positioned over right parietal regions, the limited spatial resolution of our EEG montage (60 electrodes) does not enable us to pinpoint which specific parietal regions got synchronized during right FEF rhythmic stimulation patterns. However, previous results employing diffusion imaging with participants who underwent similar non-invasive stimulation patterns and behavioral tasks (Quentin et al., 2014, 2015) suggest that fronto-parietal synchronization could operate along the 1st branch of the Superior Longitudinal Fasciculus (SLF I), part of the white matter bundle subtendng the dorsal attention network linking the FEF and the posterior intraparietal sulcus (IPS). According to well-established correlational (Corbetta et al., 2005, 2008) and causal evidence (Chanes et al., 2013; Chica et al., 2011) the network defined by the areas linked by this white matter system has been shown to play a major role in the allocation of visuo-spatial attention and the modulation of conscious perception. Accordingly, we here propose that the entrainment of oscillations within a high beta frequency (~ 30 Hz) in the right FEF could spread, presumably through white matter projections contained within the SLF I, from the FEF to right parietal regions. This specific white matter tract could underlie frequency-specific synchronization between areas of the dorsal attention orienting network responsible for enabling spatial attention, and subserve more efficient communication between frontal and parietal cortical sites. Increased coordination could be beneficial for a fast and flexible allocation of spatial attention which would enhance visual sensitivity and facilitate conscious access.

Although this outcome could be predicted on the basis of prior evidence showing a significant correlation between volume of the SLF I and TMS-induced improvements of visual perception (Quentin et al., 2014, 2015), in our ad hoc analyses, we did not entertain any strong *a priori* prediction on the loci involved in high-beta synchronization. Instead, we computed measures of synchrony between the closest electrode overlaying the right FEF and all scalp EEG leads out of a full array of 60 EEG contacts. It is hence remarkable that the statistical analyses we applied revealed a significant synchronization temporally tied to the duration of the delivered rhythmic TMS between the right FEF and electrodes over the right parietal cortex.

### Modulation of alpha oscillations

In addition to the modulation of high-beta oscillations, the implemented TMS manipulation showed effects on alpha band oscillations (Fig. 4 and 5). Since these modulations were present in active and also sham TMS conditions, we hypothesize that they did not emerge from a direct manipulation of neural activity by TMS pulses, and could instead be explained by an effect of the clicking sound accompanying the delivery of TMS pulses. The loud sound accompanying the delivery of active or sham TMS bursts, delivered in all 4 conditions 133 ms before visual target onset (in *rhythmic* or *random* bursts), could have had an alerting effect, prompting participants to engage sustained or selective attention mechanisms to focus on the task and the computer screen. The role of alpha desynchronization as a marker of attentional orienting has been well established (Capotosto et al., 2009; Klimesch et al., 1998) hence, the strong alpha desynchronization observed in all four TMS tested conditions could be consistent with this proposition.

Alternatively, the phase-locking of alpha oscillations observed after the last TMS pulse could emerge as an evoked effect of the loud TMS clicking sound, through a different mechanism. Indeed, sounds have been found to cross-modally phase-lock alpha oscillation in occipital cortices (Romei et al., 2012). Hence, an interaction between cross-modal alpha phase-locking and random TMS patterns engneered to entrain a wide range of frequencies (amongst them alpha) on different trials, could explain significantly higher alpha phase-locking over the left parietal region for active *random* compared to active *rhythmic* conditions. Much remains to be understood about these modulations in the alpha range. However, as these were not predicted, hence outside of the initial focus of this article, further studies with more adapted tasks will be required to shed more light onto these phenomena.

### Conclusions and future directions

At the instrumental or technological level, our study supports an ability to manipulate ‘at will’ inter-areal synchrony using rhythmic TMS, promising future uses of this same approach to systematically map causal synchrony interactions across frequency bands, from novel cortical sites underlying other cognitive processes. At the fundamental level, our findings extend prior evidence derived from correlational approaches and invasive electrophysiological recordings in non-human primates. To this regard, (1) they add causality to suspected associations between right frontoparietal synchronization in the high-beta band and the modulation of conscious visual performance in human participants; (2) They contribute solid evidence to support a role for high-beta right frontal oscillations and fronto-parietal synchrony as relevant physiological coding for top-down strategies to modulate visual perception; Important for the field of brain plasticity, (3) they set the stage for future applications of focal non-invasive rhythmic stimulation (via TMS or other transcranial stimulation technologies, such as tACS or ultrasound sources) to modulate local and inter-regional synchrony, to increase cognitive performance in healthy participants or improve dysfunctional synchrony patterns subtending neuropsychiatric conditions.

## Acknowledgement

Chloé Stengel was supported by a PhD fellowship from the University Pierre and Marie Curie. Marine Vernet was supported by a fellowship from the *Fondation pour la Recherche Medicale*. Julià L. Amengual was supported by a fellowship from the *Fondation Fyssen*. The activities of the laboratory of Dr. Valero-Cabré are supported by research grants IHU-A-ICM-Translationnel, ANR projet Générique OSCILOSCOPUS and Flag-era-JTC-2017 CAUSALTOMICS. The authors would also like to thank Romain Quentin, Juliette Godard and Laura Fernandez for providing help during data acquisition and the Naturalia & Biologia Foundation for financial assistance for traveling and attending meetings.

## Author contributions

Conceptualization: C.S and A.V-C. Data acquisition: M.V and A.V-C. Data analysis: C.S and J.L.A Manuscript preparation: C.S & A.V-C revised by M.V and J.LA. Supervision: A.V-C.

## Data availability statement

Data are available from the corresponding author upon request.

